# Population genomics of pneumococcal carriage in Massachusetts children following PCV-13 introduction

**DOI:** 10.1101/235192

**Authors:** PK Mitchell, T Azarian, NJ Croucher, A Callendrello, CM Thompson, SI Pelton, M Lipsitch, WP Hanage

**Author notes:** Corresponding Author: Bill Hanage,Center for Communicable Disease Dynamics,Harvard T.H. Chan School of Public Health,677 Huntington Avenue, Suite 506, Boston, MA 02115.

## Abstract

**Background:** The 13-valent pneumococcal conjugate vaccine (PCV-13) was introduced in the United States in 2010. Using a large pediatric carriage sample collected from shortly after the introduction of PCV-7 to several years after the introduction of PCV-13, we investigate alterations in the composition of the pneumococcal population following the introduction of PCV-13, evaluating the extent to which the post-vaccination non-vaccine type (NVT) population mirrors that from prior to vaccine introduction and the effect of PCV-13 on vaccine type lineages.

**Methods and Findings:** Draft genome assemblies from 736 newly sequenced and 616 previously published pneumococcal carriages isolates from children in Massachusetts between 2001 and 2014 were analyzed. Isolates were classified into one of 22 sequence clusters (SCs) on the basis of their core genome sequence. We calculated the SC diversity for each sampling period as the probability that any two randomly drawn isolates from that period belong to different SCs. The sampling period immediately after the introduction of PCV-13 (2011) was found to have higher diversity than preceding (2007) or subsequent (2014) sampling periods (Simpson’s D 2007: 0.915 95% CI [0.901, 0.929]; 2011: 0.935 [0.927, 0.942]; 2014: 0.912 [0.901, 0.923]). Amongst NVT isolates, we found the distribution of SCs in 2011 to be significantly different from that in 2007 or 2014 (Fisher’s Exact Test p=0.018, 0.0078), but did not find a difference comparing 2007 to 2014 (Fisher’s Exact Test p=0.24), indicating greater similarity between samples separated by a longer time period than between samples from closer time periods. We also found changes in the accessory gene content of the NVT population between 2007 and 2011 to have been reduced by 2014. Amongst the new serotypes targeted by PCV-13, four were present in our sample. The proportion of our sample composed of PCV-13-only vaccine serotypes 19A, 6C, and 7F decreased between 2007 and 2014, but no such reduction was seen for serotype 3. We did, however, observe differences in the genetic composition of the pre- and post-PCV-13 serotype 3 population. Our isolates were collected during discrete sampling periods from a small geographic area, which may limit the generalizability our findings.

**Conclusion:** Pneumococcal diversity increased immediately following the introduction of PCV-13, but subsequently returned to pre-vaccination levels. This is reflected in the distribution of NVT lineages, and, to a lesser extent, their accessory gene frequencies. As such, there may be a period during which the population is particularly disrupted by vaccination before returning to a more stable distribution. The persistence and shifting genetic composition of serotype 3 is a concern and warrants further investigation.

## INTRODUCTION

*Streptococcus pneumoniae* is a common bacterial colonizer of the human nasopharynx, particularly among children^1^. In Massachusetts, it has consistently been found in approximately 30% of children under the age of 7 between 2001 and 2011^2^. While colonization rarely progresses beyond asymptomatic carriage, the ubiquity of the pneumococcus leads to a substantial burden of disease, causing an estimated 4 million disease episodes, including 445,000 hospitalizations and 22,000 deaths in the United States in 2004^3^.

Conjugate vaccination has been a major advance in the reducing pneumococcal disease. The seven-valent pneumococcal conjugate vaccine (PCV-7), introduced in the United States in 2000, was highly effective in reducing overall rates of pneumococcal disease, as vaccine type (VT) pneumococci were responsible for the vast majority of cases^4-6^. Carriage of vaccine serotypes also declined, though overall carriage prevalence remained roughly constant due to serotype replacement^2,7,8^.

Despite lower overall rates of pneumococcal disease, increases were seen in the incidence of disease due to the replacement non-vaccine type (NVT) population. Serotype 19A in particular became a significant cause of invasive disease^5,9,10^. The thirteen-valent vaccine (PCV-13), introduced in 2010, extended coverage to six additional serotypes, including 19A, beyond those included in PCV-7, and has resulted in further reductions in pneumococcal disease^11^. As with PCV-7, however, overall carriage prevalence has not changed substantially^2^. Worryingly, serotype 3, a highly invasive serotype included in PCV-13, appears to have not declined as the other newly added serotypes have^2,11-14^. Given the potential for disease to arise both from replacement NVTs and persistent VTs, it remains important to monitor changes to the pneumococcal carriage population.

Pediatric pneumococcal carriage in Massachusetts has been extensively studied since shortly after the introduction of PCV-7^7^. The effects of vaccination can be seen both in the prevalence of specific lineages as well as in broader population metrics. The apparent effects of vaccination are variable depending on how the population is characterized and the timescale over which it is examined. Serotype diversity was found to have increased then stabilized following the introduction of PCV-7^15^, reflecting the selective impact of vaccines and the period while carriage replacement was taking place. Interestingly, minimal changes were found when comparing the presence and absence of specific pneumococcal genes in this population between 2001 and 2007, suggesting that the overall genetic composition of the population was not much changed other than in one of the loci conferring vaccine serotype 6B^16^. Another study considering multilocus sequence type (MLST) profiles found no significant change in diversity or population composition in the immediate aftermath of PCV-13^17^. With more time elapsed since PCV-13 introduction, it is possible to evaluate the longer-term effects of this vaccine.

Here we examine population–scale genetic changes in carriage pneumococci amongst children in Massachusetts since the introduction of PCV-13. Using genomic sequencing data for isolates collected between 2000 and 2014, we analyze alterations to the clonal composition, defined on the basis of core genome variability, and gene content of the pneumococcal NVT population following the introduction of PCV-13. Additionally, we evaluate whether serotype 3 pneumococci have declined and how they have changed through this time period.

## METHODS

### Sample Collection

Pneumococcal isolates were collected from nasopharyngeal swabs of children in Massachusetts between October and April of 2000-01, 2003-04, 2006-07, 2008-09, 2010-11 and 2013-14 as previously described^2,7,8^. Each sampling season is referred to by the later year. Pneumococcal genomes from the 2001, 2004, and 2007 sampling periods were previously published and read data for these were obtained from ENA^16^. Isolates from 2009 through 2014 were sequenced from NexteraXT genomic libraries analyzed on an Illumina MiSeq to produce paired-end 2x150 bp reads with a minimum depth of coverage of 30X.

### Genomic Processing

Draft assemblies were constructed using SPAdes v3.10 and annotated using Prokka v1.11^18,19^. Assemblies not between 1.9 and 2.3 Mb were excluded from further analysis, as were those that produced fewer than 1900 annotated coding sequences (CDS). Roary v3.10.0 was then used to identify core (present in >99% of isolates) and accessory genes and to generate a core gene alignment^20^.

### Typing

Serotype was identified using the Quellung reaction as previously described and reported for all but the 2014 sample^16,17,21^. Serotypes were checked using SRST2 v0.2.0 and a database constructed from 91 published sequences of the pneumococcal capsule biosynthetic locus^22-24^.

### Phylogenetic Analysis

The core genome alignment generated by Roary was used to construct a phylogeny using FastTree v2.1.10^25^. In order to identify clusters of related sequences (Sequence Clusters - SCs), three iterations of hierBAPS were run on the core genome alignment, setting the maximum cluster depth to 1 and maximum number of clusters to 30, 40, and 50^26^.

### Sequence Cluster Diversity

In order to determine the potential effect of PCV-13 on diversity in this population, we calculated Simpson’s D for each sampling period, for sequence clusters. This value, which represents the probability that two randomly drawn isolates from a given sampling period belong to different SCs, was calculated as 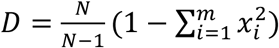, where 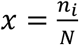, the fraction of isolates in that year belonging to sequence cluster *i* and 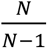 is a correction for finite sample size^27^.

Following an earlier analysis of serotype diversity in this population, Welch’s t-test was used to compare the 2007 and 2011 populations and the 2011 and 2014 populations in order to test whether SC diversity changed following the introduction of PCV-13^15^. The polyphyletic SC was excluded from these calculations.

An increase in diversity would be expected if common lineages become more rare and rare lineages become more common. To estimate the expected change in diversity we would observe if there were a smooth transition between the 2007 and 2014 population, a series of composite diversities were calculated in which the proportion belonging to each SC was a weighted combination of the 2007 and 2014 value for that SC, with the weights for the two years summing to 1. The sample size correction factor, 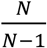, was similarly weighted.

The proportion of the population belonging to each SC and their rank order in the population were determined. As diversity increases, the shape of this distribution would be expected to flatten, with the most common lineages decreasing and the least common lineages increasing.^28^ In order to compare this distribution from the 2007 and 2014 sampling periods with that from 2011, the frequency of each SC was plotted against its rank and overlaid with the distribution from 2011. In order to determine which SCs became more or less common following the introduction of PCV13, we conducted a Fisher’s exact test for each SC comparing its frequency between the 2007 and 2014 samples.

### NVT Composition

To determine the clonal composition of the pre- and post-PCV-13 NVT population, the proportion of the NVT population belonging to each of the SCs identified by hierBAPS was calculated for 2007, 2011, and 2014. For the purpose of these analyses, serotype 6C was considered a PCV-13 type due to its cross-reactivity with serotype 6A^29,30^. Fisher’s exact test was used to determine whether these proportions varied between each pairwise combination of these three sampling periods.

We then sought to determine if the gene content of the NVT population varied between sampling periods before and after the introduction of PCV-13. Logistic models were used evaluate the extent to which individual genes became more or less common between 2007 and 2014, as well as between 2007 and 2011. Genes were excluded if they were universally present or absent in either sampling period or present or absent in fewer than 5 total isolates between the three sampling periods. For the set of genes included in both models, we calculated a linear fit comparing the regression coefficients corresponding to the time periods from 2007 to 2011 and 2007 to 2014.

In order to determine whether changes in the gene content of the NVT population from 2007 to 2011 continued, stabilized, or reversed from 2011 to 2014, we compared the observed data to hypothetical scenarios in which the 2014 population was purely reflective of the population from either the earlier sampling periods. To do this, we drew with replacement a sample of the same size as the 2014 population from either the 2007 or 2011 population. Twenty resampled populations were generated from each of 2007 and 2011, then used in place of the true 2014 population in the previous regression analyses. This process was repeated for an additional twenty resampled populations drawn from the true 2014 population in order to gauge its variability. This enabled us to evaluate the gene content of the 2014 population in relation to what would be expected if there was no overall change either from 2007 or from 2011.

### Evaluation of Serotype 3

Previous studies have noted that PCV-13 may not be as effective against serotype 3 as it is against the other serotypes included^2,11,13,14^. We compared the proportion of the pneumococcal population composed of serotype 3 between 2007, 2011, and 2014 in relation to the other three PCV-13 serotypes present in our sample, 19A, 7F, and 6C. We identified MLST profile of serotype 3 isolates using as previously described^16^. We then used RAxML to construct a phylogenetic tree based on the core genome of serotype 3 isolates to determine if the pre- and post-PCV-13 populations were genetically distnict^31^. To assess nucleotide and amino acid variation among capsular polysaccharide (CPS) loci, we mapped reads to the *S. pneumoniae* OXC141 serotype 3 reference strain (NC_017592) using SMALT v0.7.6. Single nucleotide polymorphisms (SNPs) were identified using SAMtools v1.3.1^32^. The CPS region spanning nucleotides 343,104-356,408 (*dexB* – *aliA*) was abstracted and investigated for mutations. Further, RAxML was used to construct a phylogeny of the CPS region.

## RESULTS

### Sample

A total of 1,352 isolates were included in the final analysis. The core genome consists of 1,000 genes found in at least 99% of isolates, producing an alignment 885 kb in length. A total of 10,941 genes were identified. Setting the maximum number of hierBAPS clusters to 30 and 40 produced identical results, with 21 clusters identified. With the maximum number of clusters set to 50, an additional cluster was identified and another cluster was expanded. This resulted in 22 SCs, 21 of which were monophyletic and ranged in size from 14 to 177 isolates. The other, SC1, contained 150 isolates belonging to multiple small clades or individual leaves throughout the tree and should be interpreted as containing all lineages that could not be grouped, other than on the basis of their lack of similarity to any other cluster [Fig 1].

**Figure 1:**
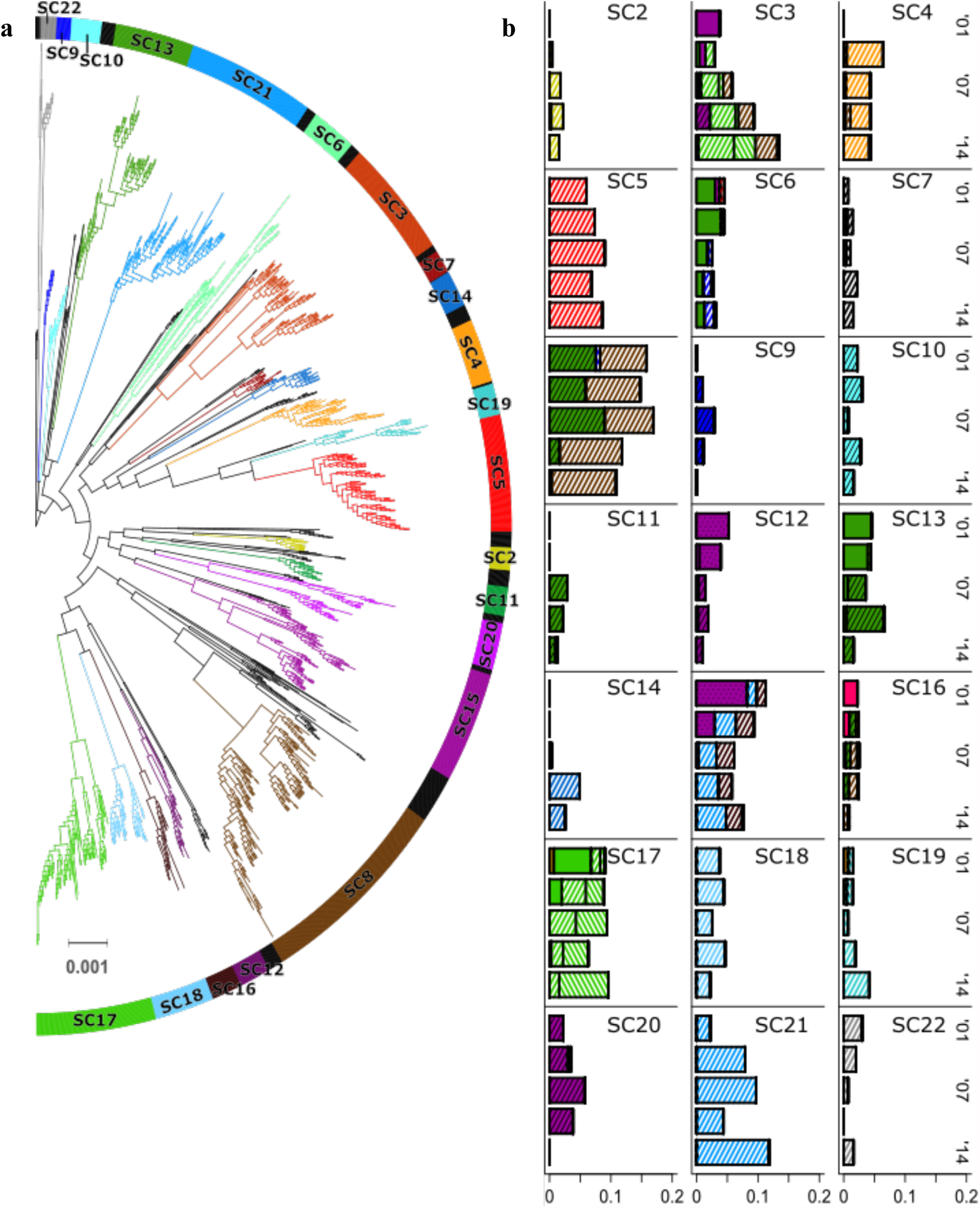
**(a)** Core genome phylogeny with SCs denoted by color. **(b)** Proportion of population in each sampling period composed of each SC, with shading indicating serotype. Solid colors are PCV-7 type, solid colors with black hatching are PCV-13, and white with colored hatching are not covered by either. Serotype 6A is dotted as it is cross-reactive with 6B, a PCV-7 type, but is itself included in PCV-13.

### Diversity

Sequence cluster diversity was calculated for each year using Simpson’s D, excluding the polyphyletic cluster SC1. Diversity was significantly higher in 2011, the first sampling period following the introduction of PCV13, than it was in either 2007 or 2014, the adjacent periods for which data were available (2007 p=0.018, 2014 p=0.00098) [Fig 2a]. A similar increase was observed after the introduction of PCV-7. The weighted diversity estimate displayed the expected increase over either the 2007 or 2014 values, but was never as high as the diversity calculated for 2011 [Fig 2b].

**Figure 2:**
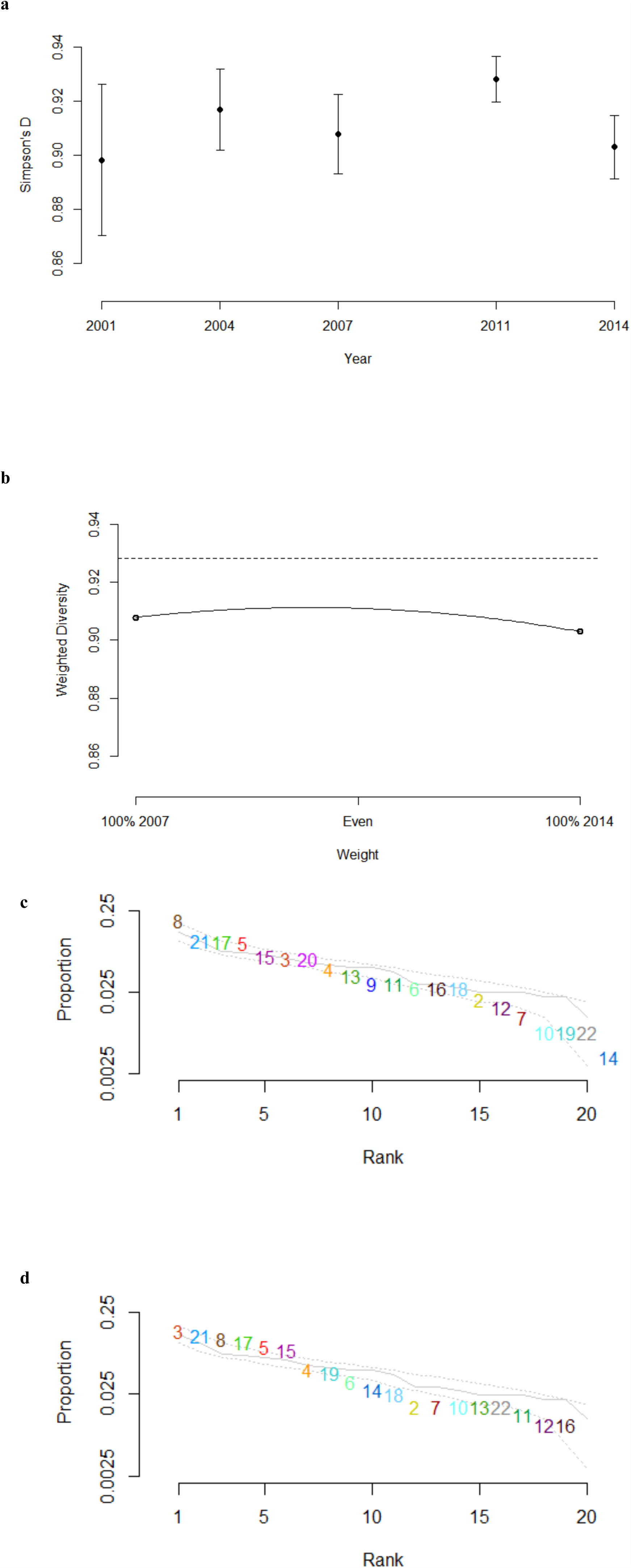
**(a)** Simpson’s diversity of SCs, excluding the polyphyletic cluster, for each sampling period. **(b)** Diversity calculated for hypothetical composites of 2007 and 2014 populations, with 2011 diversity shown as dashed line. **(c-d)** Proportion of population in **(c)** 2007 and **(d)** 2014 composed of each SC, ordered by frequency. Gray line is the corresponding distribution from 2011, with dotted lines representing 95% of values from 10000 random samples drawn from the 2011 population.

After Bonferroni correction, only 3 SCs (SC3, SC9, and SC20) changed significantly in their share of the pneumococcal population between 2007 and 2014 (Fisher’s exact test p=0.0021, 0.0022, and 4.5×10^-6^, respectively). SC3 became more common, increasing from 5.8% of the 2007 sample to 13.4% of the 2014 sample, with serogroups 23 and 15 coming to predominate over serogroup 6. Both SCs 9 and 20 are primarily composed of serotypes against which PCV-13 afforded protection (7F and 6C, respectively) and were completely absent in the 2014 sample.

The overall shape of the frequency distribution was slightly flatter in 2011 as compared to 2007 and 2014, as would be expected from the higher diversity in that sampling period. Relatively rare SCs in particular were more common in the 2011 sample than the adjacent periods [Fig 2c,d].

### NVT Composition

Non-PCV-13 types increased from 66.5% of the pneumococcal population in 2007 sampling period to 92.3% in the 2014 sampling period. Fifteen SCs had at least 1 NVT isolate. There was no significant difference between the SC distribution amongst NVTs in 2007 and 2014 (Fisher’s exact test p=0.24). There was, however, a significant difference between 2007 and 2011 (p=0.0018) and between 2011 and 2014 (p=0.0078), indicating a bounce-back effect in which the population was disrupted in 2011 but returned to its pre-vaccination state by 2014. Correspondingly, many of the common SCs that showed a distinct increase in there prevalence in the NVT population between 2007 and 2011 decreased from 2011 to 2014 while those that decreased between 2007 and 2011 increased from 2011 to 2014. [Fig 3].

**Figure 3:**
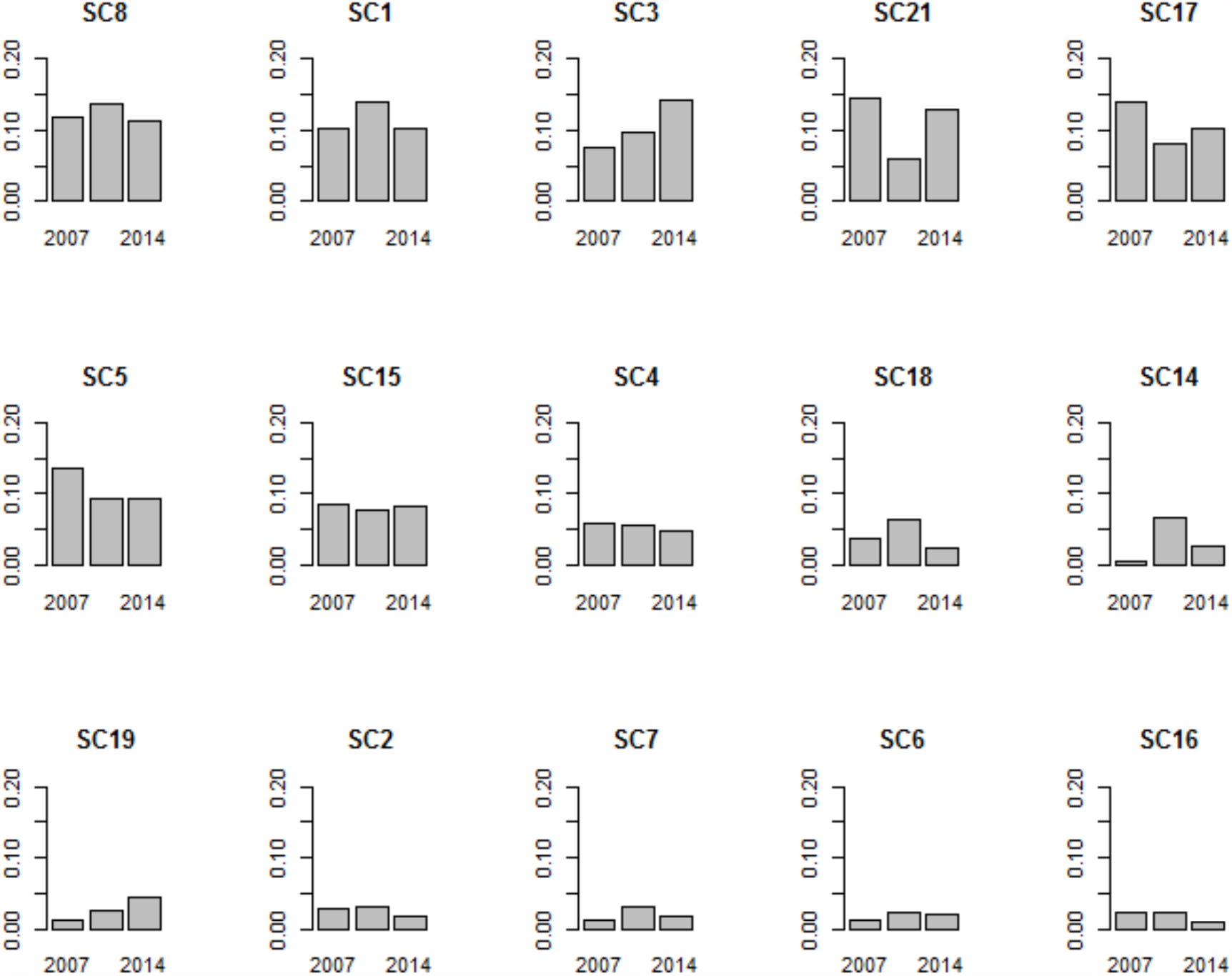
Proportion of the NVT population (i.e., those serotypes not included in PCV-13) comprised of each SC. Two additional SCs, SC11 and SC20, had a single NVT isolate and were excluded from this plot.

This bounce-back is partially reflected by the trend in gene content over time. The linear fit comparing the 2007-2011 and 2007-2014 regression coefficients for each gene had a slope of 0.62, indicating less overall change between 2007 and 2014 than between 2007 and 2011. This slope fell between those from hypothetical 2014 populations drawn from either 2007 or 2011, which clustered around a slope of 0 and 1, respectively [Fig. 4]. This indicates that while the direction in which genes changed in frequency from 2007 to 2011 was generally preserved through 2014, the trend was partially counteracted between 2011 and 2014 with genes returning closer to their 2007 levels prior to the introduction of PCV-13.

**Figure 4:**
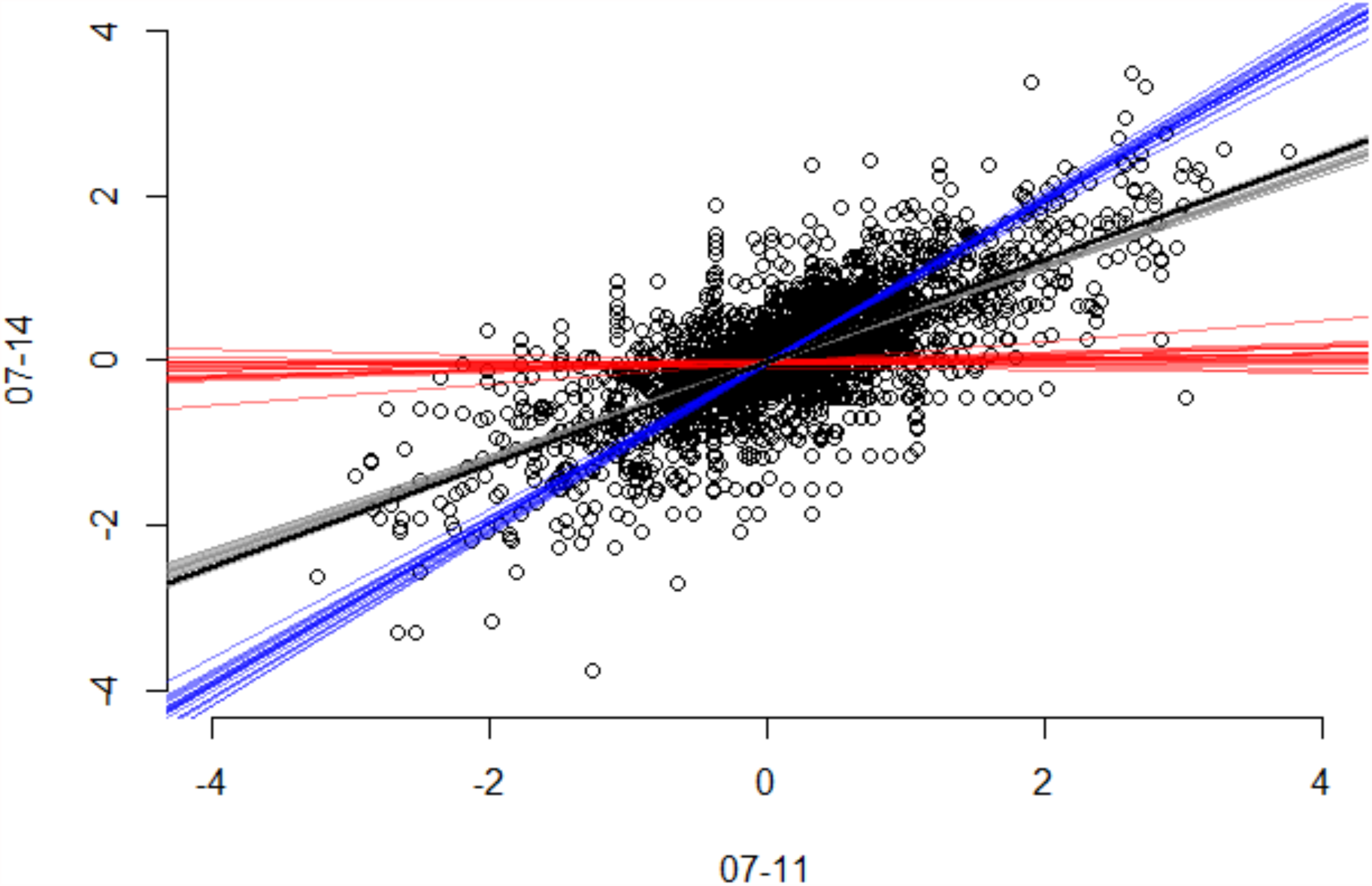
Regression coefficients comparing gene content of the NVT population from 2007 to 2011 and 2014. Black circles correspond to the coefficients with individual genes, with a linear fit to the data shown in black. Fits in which a hypothetical 2014 population was drawn from either the 2007, 2011, or 2014 population are shown in red, blue, and gray, respectively.

### Persistence of Serotype 3

In order to evaluate whether the whether the new serotypes included in PCV-13 decreased following its introduction, we conducted a Fisher’s exact test comparing the 2007 and 2014 carriage share of serotypes 19A, 6C, 7F, and 3. While serotypes 19A, 6C and 7F all showed significant reductions between the two time periods (p<0.0001, p=0.00014, p=0.0011, respectively), serotype 3 had no such change (p=0.46) [Fig 5].

**Figure 5:**
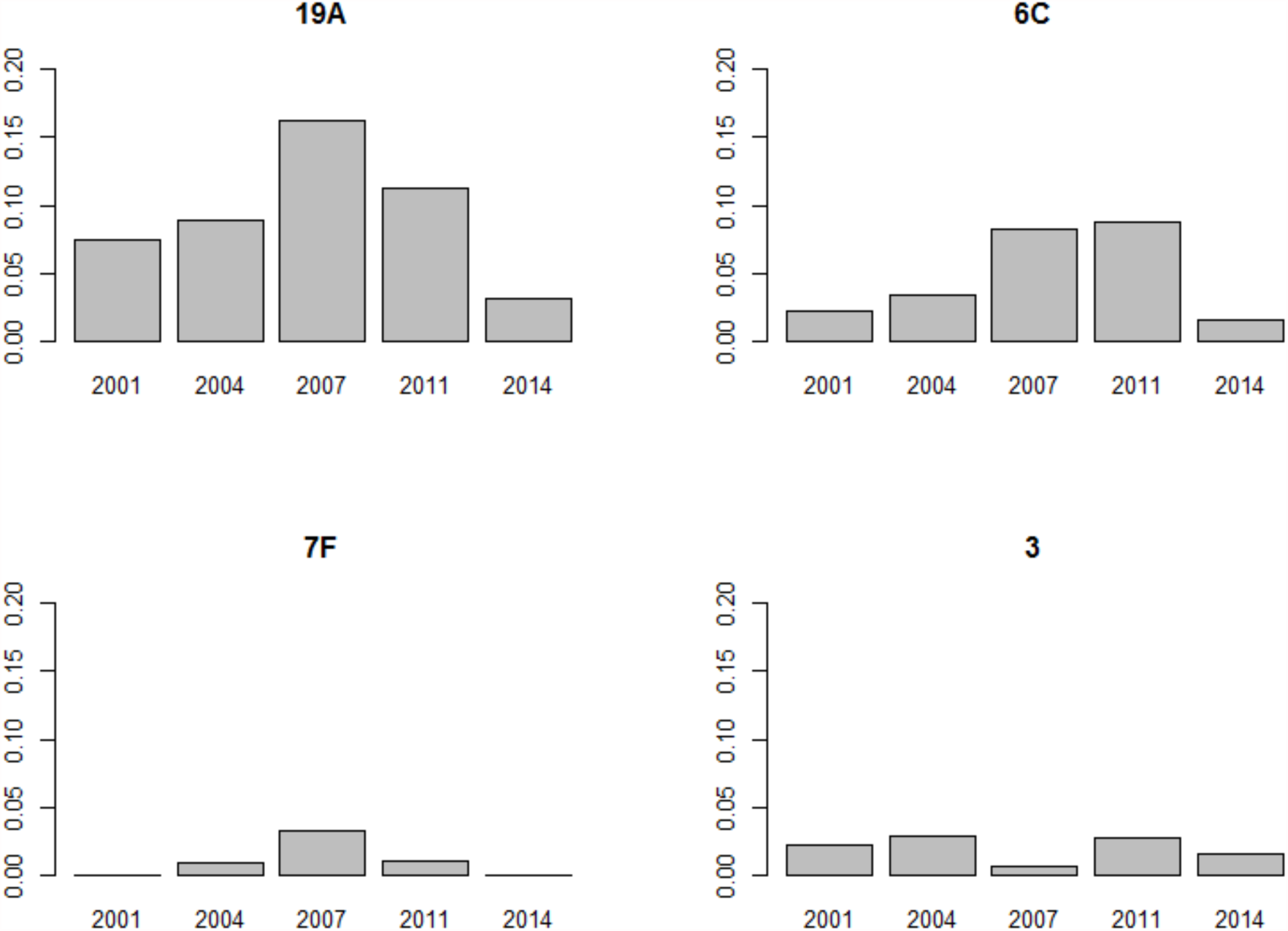
Proportion of the population in each sampling period comprised of the serotypes included in PCV-13 but not PCV-7. As a note, PCV-7 was introduced in the United States in 2000 and PCV-13 was introduced in 2010.

To test if the persistence of serotype 3 may be related to some genetic factor, we assessed population structure and CPS nucleotide variation. All of the serotype 3 isolates clustered in the same SC and were MLST sequence type (ST) 180 belonging to the Netherlands^3^–31 (PMEN31) clone CC180. While all isolates clustered into the same SC, there was a distinct bifurcation in the phylogeny [Fig 6]. Of the 28 serotype 3 isolates, 16 fell into one subclade and 12 into the other. In the larger subclade, 4 isolates (25%) are from 2011 or 2014, after the introduction of PCV-13. The other subclade contains 11 (92%) post-PCV-13 isolates (χ^2^ p=0.0018). Further assessment of CPS showed low nucleotide diversity [mean pairwise SNP distance:1.5 (S.E. 0.7)] and only four polymorphic amino acids, none of which segregated the subclades. However, the CPS phylogeny recapitulated the bifurcation in the core genome phylogeny, with all isolates belonging to the post-PCV-13 subclade displaying as highly clustered.

**Figure 6:**
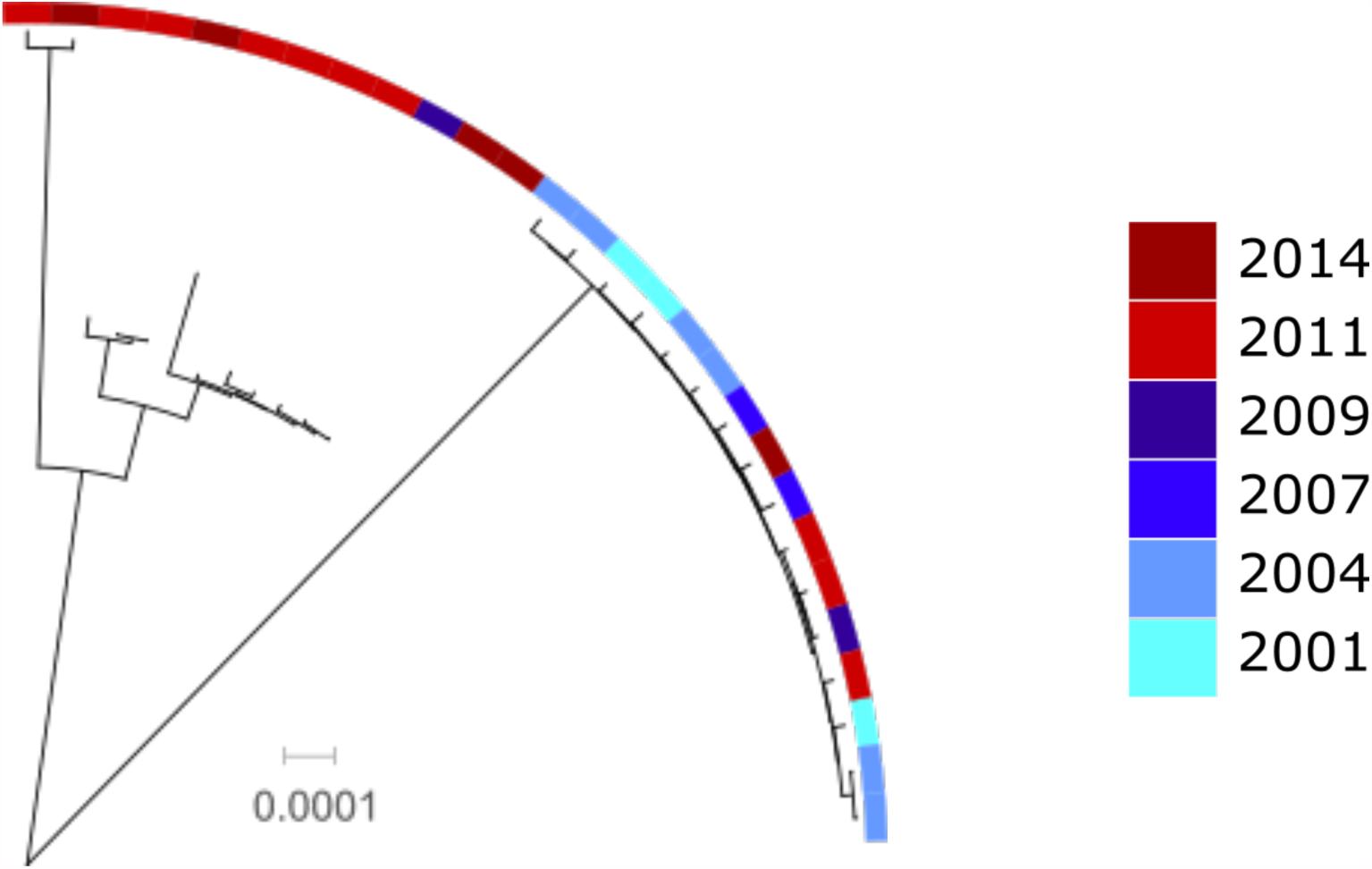
Serotype 3 phylogeny, with sampling period shown by color. Isolates collected before the introduction of PCV13, shown in blue, are found primarily on one monophyletic clade of the tree, while post-introduction isolates, indicated by red, are primarily on the other.

## DISCUSSION

Here we have analyzed a sample of carriage pneumococci collected in Massachusetts between the winters of 2000-01 and 2013-14, focusing primarily on changes occurring following the introduction of PCV-13. Using genomic data, we find that the NVT population in the most recent sampling period more closely reflects that of our last full pre-PCV13 sample than our first post PCV-13 sample. This suggests a return to equilibrium following disruption by vaccine, which is consistent with observations made following the introduction of PCV-7 in the same population^33^, but now with the added resolution offered by genomic data. We also find that serotype 3 CC180 has been more persistent than other serotypes added for PCV-13, but a different subclade of this lineage now predominates.

Given the value of being able to predict the composition of the pneumococcal population following PCV use, the pattern observed amongst the NVTs is quite interesting. Our 2014 sample appears to be broadly a reflection of the 2007 sample, but 2011 is unlike either. As such, it is possible that the pre-vaccine NVT population may be a good predictor of the post-vaccine population, but that the disruption caused by vaccine introduction can temporarily interrupt this pattern. Some of this could be due to variation in the age of children who have been vaccinated, which should increase over time as vaccinated children age. The observed increase in SC diversity in the immediate post-vaccine period, with the most common lineages making up a smaller proportion of the total population, may provide an enhanced opportunity for rarer lineages to increase. Considering this scenario, lineages such as SCs 3, 14, and 19 (serotypes 23A/15BC, 21 and 33F, respectively), may have a similar trajectory to that of serotype 19A ST320 after pcv-7^5,9,10,34^. It has also recently been suggested that negative frequency dependent selection on elements of the accessory genome could be responsible for structuring the pneumococcal population at both spatial and temporal scales^35^. Further observation will help determine the role of this and whether these or other lineages become more substantial contributors to both carriage and invasive disease.

Previous studies have indicated that PCV-13 may not be as effective against serotype 3 as it is against other included serotypes^2,11,13,14^. The shift we observed in the serotype 3 CC180 population following the introduction of PCV-13 may reflect a similar phenomenon to that leading to the recognition of serotype 6C as distinct from 6A following the introduction of PCV-7^36,37^. The dominant lineage pre-PCV-13 was also more homogenous (i.e., less diverse) than the post-vaccination population, so it is possible that the immunity generated against serotype 3 by PCV-13 is narrowly tailored to that subset of the population. At present, little genetic variation among the CPS loci was observed, suggesting an alternative explanation for the recent post-PCV-13 emergent subclade. Given its propensity for causing disease, the persistence of serotype 3 despite its inclusion in PCV-13 warrants further investigation.

The response of the pneumococcal population to serotype-targeting conjugate vaccines may also provide insights for other pathogens for which vaccines have been targeted at or differentially affect a subset of their population. The efficacy of the RTS,S malaria vaccine appears to be partially dependent on how well the circumsporozoite protein of a given *Plasmodium* type matches that in the vaccine^38^. There has also been interest in understanding how the strain dynamics and epidemiology of meningococcal disease caused by the bacteria *Neisseria meningitidis* will be affected by the rollout of vaccinations against a variety of serogroups^39,40^. While each of these disease systems is different, there is some potential for findings in one to inform hypotheses for how others will behave.

Pneumococcal epidemiology has changed substantially as a result of conjugate vaccination. While PCVs have been highly effective in reducing the incidence of pneumococcal disease^4,5,11^, continued vigilance is necessary to monitor for, and respond to, the emergence of potentially dangerous lineages not protected against by current vaccine formulations.

## References

Mehr S, Wood N. Streptococcus pneumoniae – a review of carriage, infection, serotype replacement and vaccination. Paediatr Respir Rev. January 2012:2–8. doi:10.1016/j.prrv.2011.12.001.

Lee GM, Kleinman K, Pelton SI, et al. Impact of 13-valent pneumococcal conjugate vaccination on Streptococcus pneumoniae carriage in young children in Massachusetts. J Pediatric Infect Dis Soc. 2014;3(1):23–32. doi: 10.1093/jpids/pit057.

Huang SS, Johnson KM, Ray GT, et al. Healthcare utilization and cost of pneumococcal disease in the United States. Vaccine. 2011;29(18):3398–3412. doi:10.1016/j.vaccine.2011.02.088.

Whitney CG, Farley MM, Hadler J, et al. Decline in Invasive Pneumococal Disease after the Introduction of Protein-Polysachharide Conjugate Vaccine. N Engl J Med. 2003;348(18):1737–1746.

Pilishvili T, Lexau C, Farley MM, et al. Sustained reductions in invasive pneumococcal disease in the era of conjugate vaccine. J Infect Dis. 2010;201(1):32–41. doi:10.1086/648593.

Hausdorff WP, Bryant J, Paradiso PR, Siber GR. Which pneumococcal serogroups cause the most invasive disease: implications for conjugate vaccine formulation and use, part I. Clin Infect Dis. 2000;30(1):100–121. doi:10.1086/313608.

Huang SS, Platt R, Rifas-Shiman SL, Pelton SI, Goldmann D, Finkelstein J a. Post-PCV7 changes in colonizing pneumococcal serotypes in 16 Massachusetts communities, 2001 and 2004. Pediatrics. 2005;116(3):e408-13. doi:10.1542/peds.2004-2338.

Huang SS, Hinrichsen VL, Stevenson AE, et al. Continued Impact of Pneumococcal Conjugate Vaccine on Carriage in Young Children. Pediatrics. 2009;124(1):e1-e11. doi:10.1542/peds.2008-3099.

Moore MR, Gertz RE, Woodbury RL, et al. Population snapshot of emergent Streptococcus pneumoniae serotype 19A in the United States, 2005. JInfect Dis. 2008;197(7):1016–1027. doi:10.1086/528996.

Pelton SI, Huot H, Finkelstein J a, et al. Emergence of 19A as virulent and multidrug resistant Pneumococcus in Massachusetts following universal immunization of infants with pneumococcal conjugate vaccine. Pediatr Infect Dis J. 2007;26(6):468–472. doi:10.1097/INF.0b013e31803df9ca.

Moore MR, Link-Gelles R, Schaffner W, et al. Effect of use of 13-valent pneumococcal conjugate vaccine in children on invasive pneumococcal disease in children and adults in the USA: analysis of multisite, population-based surveillance. Lancet Infect Dis. 2015; 15(3):301–309. doi:10.1016/S1473-3099(14)71081-3.

Yildirim I, Hanage WP, Lipsitch M, et al. Serotype specific invasive capacity and persistent reduction in invasive pneumococcal disease. Vaccine. 2010;29(2):283–288. doi:10.1016/j.vaccine.2010.10.032.

Andrews NJ, Waight PA, Burbidge P, et al. Serotype-specific effectiveness and correlates of protection for the 13-valent pneumococcal conjugate vaccine: a postlicensure indirect cohort study. Lancet Infect Dis. 2014;14(9):839–846. doi:10.1016/S1473-3099(14)70822-9.

Harboe Z, Dalby T. Impact of 13-Valent Pneumococcal Conjugate Vaccination in Invasive Pneumococcal Disease Incidence and Mortality. Clin Infect Dis. 2014;59:1066–1073. doi:10.1093/cid/ciu524.

Hanage WP, Finkelstein JA, Huang SS, et al. Evidence that pneumococcal serotype replacement in Massachusetts following conjugate vaccination is now complete. Epidemics. 2010;2(2):80–84. doi:10.1016/j.epidem.2010.03.005.

Croucher NJ, Finkelstein JA, Pelton SI, et al. Population genomics of post-vaccine changes in pneumococcal epidemiology. Nat Genet. 2013;45(6):656–663. doi :10.1038/ng.2625.

Chang Q, Stevenson AE, Croucher NJ, et al. Stability of the pneumococcal population structure in Massachusetts as PCV13 was introduced. BMC Infect Dis. 2015;15:68. doi:10.1186/s 12879-015-0797-z.

Bankevich A, Nurk S, Antipov D, et al. SPAdes: a new genome assembly algorithm and its applications to single-cell sequencing. J Comput Biol. 2012;19(5):455–477. doi:10.1089/cmb.2012.0021.

Seemann T. Prokka: rapid prokaryotic genome annotation. Bioinformatics. 2014;30(14):2068–2069. doi:10.1093/bioinformatics/btu153.

Page AJ, Cummins CA, Hunt M, et al. Roary: Rapid large-scale prokaryote pan genome analysis. Bioinformatics. 2015;31(22):btv421. doi:10.1093/bioinformatics/btv421.

Hanage WP, Bishop CJ, Huang SS, et al. Carried pneumococci in Massachusetts children: the contribution of clonal expansion and serotype switching. Pediatr Infect Dis J. 2011;30(4):302–308. doi:10.1097/INF.0b013e318201a154.

Inouye M, Dashnow H, Raven L-A, et al. SRST2: Rapid genomic surveillance for public health and hospital microbiology labs. Genome Med. 2014;6(11):90. doi:10.1186/s13073-014-0090-6.

Bentley SD, Aanensen DM, Mavroidi A, et al. Genetic analysis of the capsular biosynthetic locus from all 90 pneumococcal serotypes. PLoS Genet. 2006;2(3):e31. doi:10.1371/journal.pgen.0020031.

Park IH, Park S, Hollingshead SK, Nahm MH. Genetic basis for the new pneumococcal serotype, 6C. Infect Immun. 2007;75(9):4482–4489. doi:10.1128/IAI.00510-07.

Price MN, Dehal PS, Arkin AP. FastTree: Computing Large Minimum Evolution Trees with Profiles instead of a Distance Matrix. Mol Biol Evol. 2009;26(7):1641–1650. doi:10.1093/molbev/msp077.

Cheng L, Connor TR, Sirén J, Aanensen DM, Corander J. Hierarchical and spatially explicit clustering of DNA sequences with BAPS software. Mol Biol Evol. 2013;30(5):1224–1228. doi:10.1093/molbev/mst028.

Simpson EH. Measurement of Diversity. Nature. 1949;163:688–688. doi:10.1038/163688a0.

Dagan R, Patterson S, Juergens C, et al. Comparative Immunogenicity and Efficacy of 13-Valent and 7-Valent Pneumococcal Conjugate Vaccines in Reducing Nasopharyngeal Colonization: A Randomized Double-Blind Trial. Clin Infect Dis. 2013;57(7):952–962. doi: 10.1093/cid/cit428.

Cooper D, Yu X, Sidhu M, Nahm MH, Fernsten P, Jansen KU. The 13-valent pneumococcal conjugate vaccine (PCV13) elicits cross-functional opsonophagocytic killing responses in humans to Streptococcus pneumoniae serotypes 6C and 7A. Vaccine. 2011;29(41):7207–7211. doi:10.1016/j.vaccine.2011.06.056.

Stamatakis A. RAxML-VI-HPC: Maximum likelihood-based phylogenetic analyses with thousands of taxa and mixed models. Bioinformatics. 2006;22(21):2688–2690. doi:10.1093/bioinformatics/btl446.

Li H, Handsaker B, Wysoker A, et al. The Sequence Alignment/Map format and SAMtools. Bioinformatics. 2009;25(16):2078–2079. doi:10.1093/bioinformatics/btp352.

Hanage WP, Fraser C, Tang J, Connor TR, Corander J. Hyper-Recombination, Diversity, and Antibiotic Resistance in Pneumococcus. Science (80-). 2009;324(5933):1454–1457. doi:10.1126/science.1171908.

Hanage WP, Bishop CJ, Lee GM, et al. Clonal replacement among 19A Streptococcus pneumoniae in Massachusetts, prior to 13 valent conjugate vaccination. Vaccine. 2011;29(48):8877–8881. doi:10.1016/j.vaccine.2011.09.075.

Jukka Corander, Christophe Fraser, Michael U. Gutmann, Brian Arnold, William P. Hanage, Stephen D. Bentley, Marc Lipsitch NJC. Frequency-dependent selection in vaccine-associated pneumococcal population dynamics. Nat Ecol Evol. October 2017:In press. doi:10.1038/s41559-017-0337-x.

Park IH, Moore MR, Treanor JJ, et al. Differential effects of pneumococcal vaccines against serotypes 6A and 6C. J Infect Dis. 2008;198:1818–1822. doi:10.1086/593339.

Park IH, Pritchard DG, Cartee R, Brandao A, Brandileone MCC, Nahm MH. Discovery of a new capsular serotype (6C) within serogroup 6 of Streptococcus pneumoniae. J Clin Microbiol. 2007;45(4):1225–1233. doi:10.1128/JCM.02199-06.

Neafsey DE, Juraska M, Bedford T, et al. Genetic Diversity and Protective Efficacy of the RTS,S/AS01 Malaria Vaccine. N Engl J Med. 2015;373(21):2025–2037. doi:10.1056/NEJMoa1505819.

Halperin SA, Bettinger JA, Greenwood B, et al. The changing and dynamic epidemiology of meningococcal disease. Vaccine. 2012;30:26–36. doi:10.1016/j.vaccine.2011.12.032.

Ali O, Aseffa A, Bedru A, et al. The diversity of meningococcal carriage across the African meningitis belt and the impact of vaccination with a group a meningococcal conjugate vaccine. J Infect Dis. 2015;212:1298–1307. doi:10.1093/infdis/jiv211.

